# A neurocomputational model of human frontopolar cortex function

**DOI:** 10.1101/037150

**Authors:** Alexandre Hyafil, Etienne Koechlin

## Abstract

The frontopolar cortex (FPC), the most anterior part of the lateral prefrontal cortex corresponding to Brodmann’s area 10, is involved in human high-order cognition, including reasoning, problem-solving and multitasking. Its specific contribution to prefrontal executive function, however, remains unclear. A neurocomputational model suggests that the FPC implements a basic process referred to as cognitive branching that maintains a task in a pending state during the execution of another, and enables to revert back to it upon completion of the ongoing one. However, the FPC is engaged in other cognitive functions including prospective memory, relational reasoning, episodic memory retrieval and attentional set-shifting, which are not directly linked to the notion of cognitive branching. Here we used a neurocomputional branching model to simulate the involvement of the FPC in these various cognitive functions. Simulation results indicate that the model accounts for the variety of FPC activations observed in these various experimental paradigms. Thus, the present study provides theoretical evidence suggesting that all these behavioral paradigms implicitly involve branching processes, and supports the idea that cognitive branching is the core function of the human frontopolar cortex.

## Introduction

The most anterior part of the lateral prefrontal cortex, the so-called frontopolar cortex (FPC) corresponds to a cyto-architectonically defined brain region (Brodmann’s area 10) that develops lately, both phylogenetically and ontogenically (Semendeferi K et al., 2001). The FPC forms the apex of a functional hierarchy of control processes originating from the premotor cortex and underlying executive control, i.e. the coordination of actions and thoughts in relation with internal goals (Badre D and M D'Esposito, 2007; Koechlin E et al., 2003; Koechlin E and C Summerfield, 2007) (Miller EK and JD Cohen, 2001). The specific contribution of the frontopolar cortex to executive control, however, remains poorly understood, especially since patients with frontopolar lesions often show little impairments in standardized neuropsychological prefrontal tests (Burgess PW, 2000). Based on recent results from neuroimaging studies, several hypotheses have been put forward. In particular, the FPC was alternatively proposed to subserve cognitive branching (i.e. postponing the execution of a primary task until completion of a secondary task)(Koechlin E et al., 1999), to control internally generated thoughts (Burgess PW et al., 2005; Christoff K et al., 2003), to integrate outcomes of multiple tasks (Ramnani N and AM Owen, 2004) and to assist in exogenous attentional set-shifting (Pollmann S, 2001). These accounts, however, have often been formulated on the basis of elusive concepts, so that it remains largely unclear how they relate to each other and, more problematically, whether these hypotheses can account for the engagement of the FPC throughout a variety of behavioral paradigms (Burgess PW *et al.*, 2005; Christoff K and JD Gabrieli, 2000; Ramnani N and AM Owen, 2004) involving multitasking, prospective memory (Burgess PW et al., 2001), relational reasoning paradigms (Bunge SA et al., 2004; Christoff K et al., 2001), attentional set-shifting (Pollmann S, 2004) and episodic memory retrieval (Buckner RL and ME Wheeler, 2001).

To clarify the role of the frontopolar cortex in executive control, we recently developped a neurocomputational model that describes how FPC function may emerge from interactions with neighboring prefrontal regions constituting its major input/output pathways: namely the lateral and the orbital/medial prefrontal cortex, implicated in cognitive and motivational control, respectively (Koechlin 2007). More specifically, the model suggests that the FPC selectively implements branching processes: based on motivational signals (i.e. expected future rewards associated with distinct tasks) originating from the orbital/medial prefrontal cortex, the FPC temporarily maintains the second most rewarding task in a pending state, while the lateral prefrontal cortex controls the execution of the most rewarding task. The mechanism enables the execution of the pending task upon completion of the ongoing one, even without any explicit retrieval cues. Such a cognitive branching mechanism appears critical for overcoming the serial constraint that bears upon executive control. Indeed, previous behavioral studies revealed that the central executive system is unable to control the execution of multiple tasks at one time (Pashler H, 2000; Sigman M and S Dehaene, 2005). Moreover, neuropsychological studies revealed that multitasking behaviors are specifically impaired in patients with frontopolar lesions (Burgess PW et al., 2000), while neuroimaging studies on healthy subjects showed that multitasking robustly engage the FPC (Braver TS and SR Bongiolatti, 2002; Koechlin E *et al.*, 1999; Koechlin E et al., 2000).

A remaining issue, however, is whether this cognitive branching model accounts for the involvement of the FPC in the wide variety of behavioral paradigms described above that are not intuitively linked to multitasking. Here, we present a brief overview of the branching model proposed by (Koechlin 2007) and we report results from model simulations mimicking subjects’ performance in these behavioral paradigms.

## Model overview

### Architecture and connectivity

The model implements an executive system that serially organizes *task-set* execution. We refer to the term *task-set* as an association combining behavioral rules, external cues that trigger implementation of these rules, and rewards expected from their execution. Such associations are activated in response to external cues and maintained over time to enforce specific behavioral rules for as long as their associated expected rewards remain valuable enough (Miller EK and JD Cohen, 2001). The core property of the model is to decide and allow the maintenance of a task-set in a pending state during the execution of another.

The model describes functional interactions between the frontopolar cortex and two neighboring prefrontal regions, the orbital/medial and lateral prefrontal cortex. Both cortical regions share important reciprocal connections with the frontopolar cortex and presumably constitute the major input-output system of the frontopolar cortex (Barbas H and DN Pandya, 1989; Petrides M and DN Pandya, 1999). The model aims to capture the key computational features of frontopolar function, and thus represents an oversimplification of underlying neuronal mechanisms. However, the model is consistent with basic properties of neuronal processing.

The model is composed of three interconnected modules referred to as LPC, FPC and OFC representing the lateral, frontopolar and orbital/medial prefrontal regions, respectively (Fig. 1). Each module includes interconnected processing units modeling distinct populations of neurons. In agreement with previous studies on the specific contribution of each prefrontal sector to executive control (Matsumoto K and K Tanaka, 2004; Miller EK and JD Cohen, 2001; Rolls ET, 2004), we assumed that LPC and OFC code for ongoing behavioral rules and expected rewards, respectively. Cross-module connections form closed loop circuits, so that projections across modules mainly connect units related to the same task-set.

**Figure 1.**
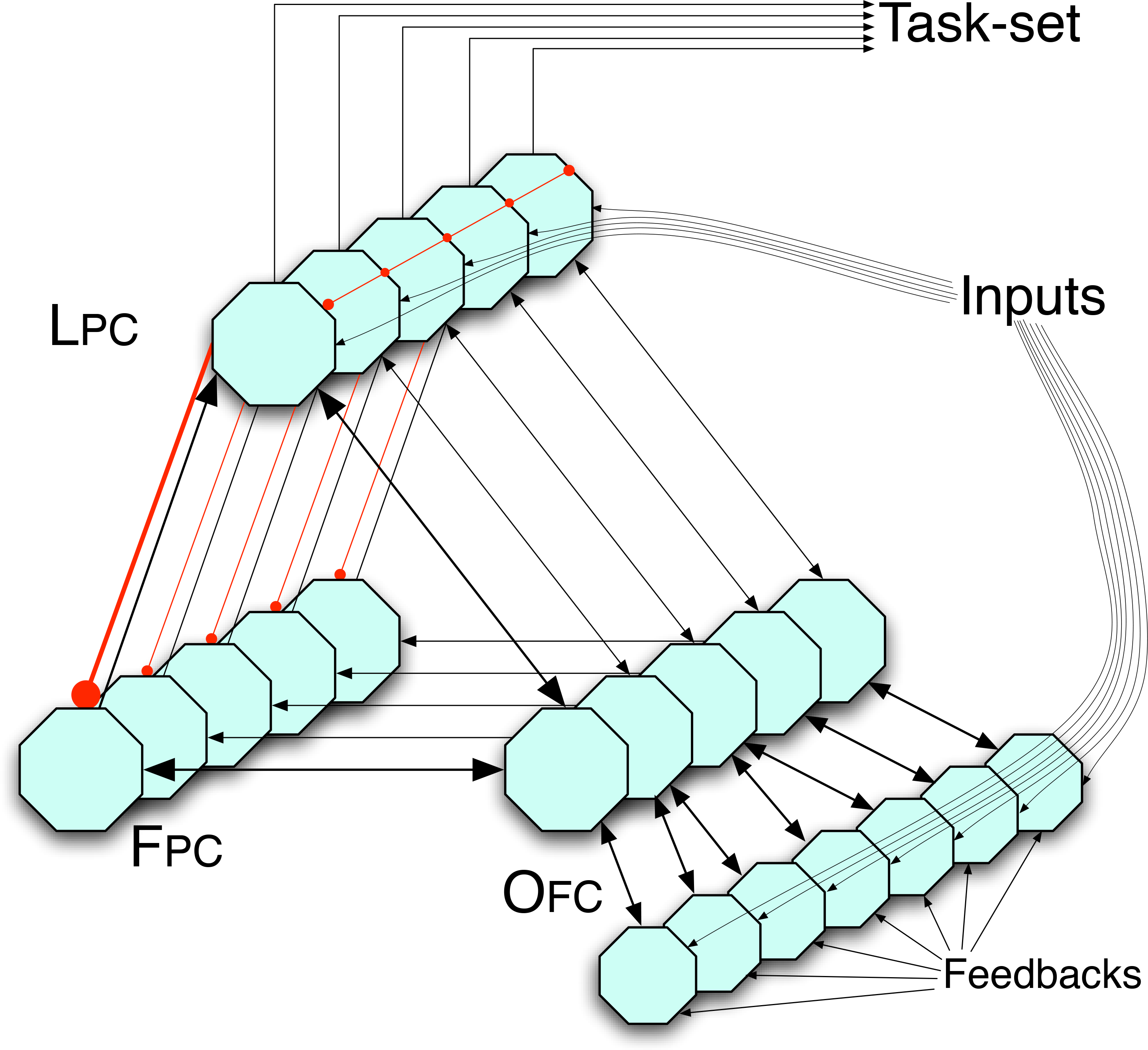
Architecture of the model. Octagons represent LPC, OFC and FPC units, representing distinct neural populations within these prefrontal areas. Black arrows and red arrows with round head indicate excitatory and inhibitory interactions respectively. Strong lateral inhibition and auto-excitation occur in LPC and FPC, implementing winner-take-all competition in both modules. External cues provide inputs to associated units in both LPC and OFC input layer. The weight of the projection from a cue-specific OFC input to a task-set specific OFC reward unit encodes the value of rewards expected from performing the related task-set, as predicted by the associated external cue. Sustained activation of LPC units are assumed to reflect ongoing performance of the associated task-set, through top-down interactions with more posterior brain areas; neural processing in these posterior areas were not modeled. During task-set performance, OFC receives additional inputs that update the encoded value of expected future rewards. This mechanism allows to clear all activity in the network related to a task-set when that task-set is completed.

Active units in LPC represent the ongoing task-set: top-down projections from LPC units to posterior prefrontal areas are assumed to select the behavioral rules associated with this task-set. For simplicity, we did not model the implementation of these rules in posterior regions. Each LPC unit receives bottom-up excitatory projections from posterior associative areas signaling the occurrence of external cues associated with the corresponding task-set (e.g. instruction cues). Strong lateral inhibition across LPC units implement winner-take-all mechanisms enforcing a single task-set to be performed at one time. The ongoing task-set is maintained through self-excitation within LPC and reciprocal excitatory connections between LPC and OFC units.

OFC units activity encodes expected reward values associated with each task-set, i.e. the net total amount of future reward expected from task-set performance. OFC units receive external signals through an *input layer.* Feedforward connections from that input layer to OFC reward layer store the value of rewards expected from performing the related task-set, given external inputs. OFC reward units continuously integrate these inputs to update the value of expected future rewards associated with each task-set; such reward evaluations provide a bias for task-set selection in LPC. Reciprocal excitatory projections from LPC to OFC reward units enable the maintenance of these activities after the offset of external cues. Thus, in response to external inputs to both modules, reciprocal excitatory interactions between LPC and OFC enable selection and maintenance of the task-set associated with the largest expected reward. OFC unit activity ultimately vanishes when no more reward is expected from the selected task-set, which may correspond to task-set completion. Consistently, activity of the associated LPC unit vanishes too.

FPC provides a temporary storage device required for branching, i.e. maintaining pending task-sets for future performance and preventing interference with the ongoing taskset represented in LPC. Unlike LPC and OFC units, FPC units receive no external inputs and interact with LPC and OFC modules only. Reciprocal excitatory connections between OFC and FPC and lateral inhibition within FPC allow FPC to select and maintain task-sets associated with large enough expected rewards. Lateral inhibition between OFC units is weak enough to enable OFC to simultaneously represent the expected rewards associated with distinct task-sets, namely those selected in LPC and FPC. Critically, the model assumes asymmetrical interactions between LPC and FPC. Bottom-up inhibitory projections from LPC to FPC prevent FPC from encoding the same task-set than the one selected in LPC. Conversely, top-down excitatory projections from FPC to LPC enable the pending task-set encoded in FPC to “migrate” to LPC as soon as the expected future rewards associated with the pending task-set become larger than those associated with the ongoing task-set encoded in LPC (e.g. when the ongoing task-set is completed).

Thus, a key feature of the model is the inhibitory interaction from LPC to FPC units, which is required for FPC units to encode task-sets distinct from those encoded in LPC. This assumption may appear inconsistent with the fact that long-distance corticocortical connections are excitatory (white & Keller 1989, dans Barbas sept 2005). However, as previously reported (Barbas et al.), the influence of long-distance connections on local neuronal processing may instead be inhibitory when connection targets are inhibitory neurons.

### Unit activity equations

Unit activity in all three modules are modeled using the classical Wilson & Cowan’s model (Wilson HR and JD Cowan, 1973):

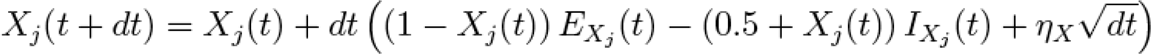

where *X_j_(t)* denotes the activity of unit *j* in module *X* (being LPC, OFC or FPC) at time *t, Ex_j_(t)* and *Ix_j_(t)* excitatory and inhibitory interactions, respectively. *η_x_* denotes the standard deviation of normally distributed noise within module *X* (with the constraint that unit activity remains positive). Excitation and inhibition terms in each module are derived from the pattern of connectivity described above:

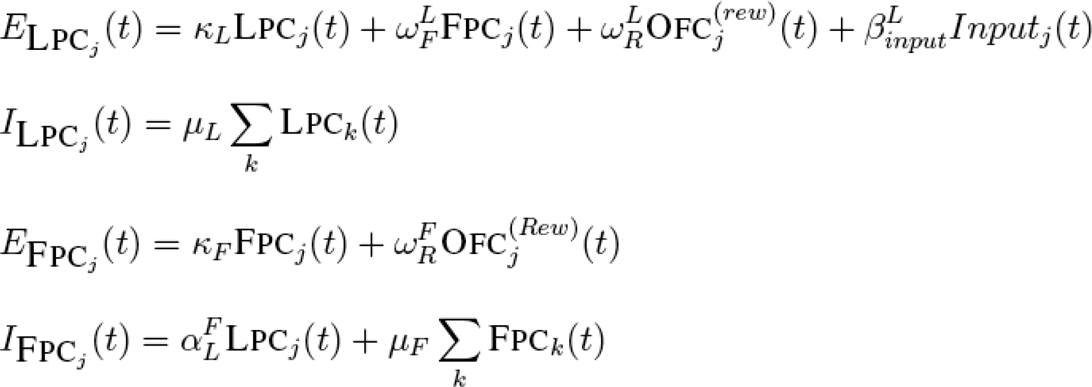

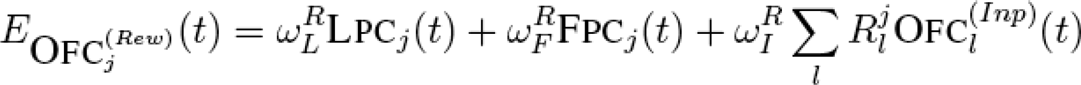

where R^J^_i_ denotes the value of the reward expected from execution of task-set *j* given the occurrence of cue l. Indices *Inp* and *Rew* refer to input and reward layers in O_FC_. Index *k* lists all units in each module, except for OFC input layer where index *l* was used.

Activation rules of processing units in LPC and FPC include standard linear summation of inputs, including self-excitation, lateral inhibition between processing units, feedforward and feedback long-distance excitation and inhibition across cortical areas. The model does not include lateral excitation, as the processing units used in the model describe populations of neighboring neurons that share excitatory reciprocal connections rather than single neurons.

OFC unit activity follows an activation rule that reproduces the known properties of neurons in the medial/orbital prefrontal regions, namely neuronal activity reflects ongoing evaluation of expected future reward based on learned associations between external signals and rewards (Rolls ET, 2004).

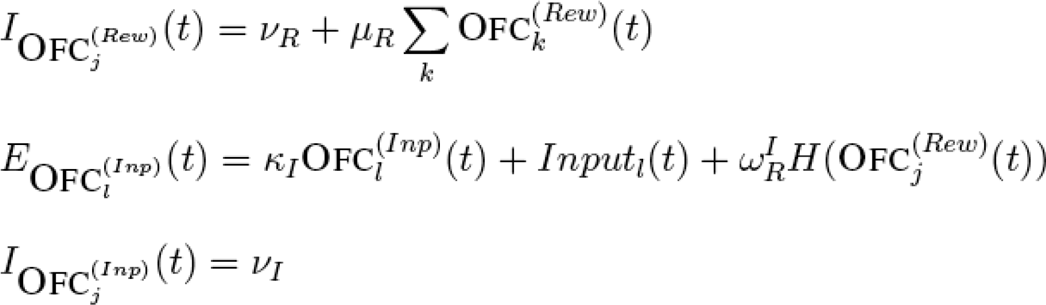

where *H(x)* is the step function worth 1 if x > .5 and 0 otherwise.

Finally, it is worth noting that according to these activation rules, LPC and FPC units exhibit modulations of activity by expected rewards through excitatory projections from OFC units. Consistently, neuronal activity in monkeys as well as activations in lateral prefrontal and frontopolar regions in human neuroimaging studies were found to be sensitive to reward expectations (Leon MI and MN Shadlen, 1999; Pochon JB et al., 2002).

### Simulation parameters

For the simulations, we used the following set of parameters: КL = КF = КI = 0.5, ω^L^F = 0.1, ω^R^L = 0.1, ω^R^F = 0.055, ω^L^R= 0.6, ω^F^R= 1.7, ωV= 0.08, ω^R^i = 0.05, a^F^L= 5, ß^L^input = 0. 1, μ_*L*_ = μ_F_ = 1, μ_R_ = 0.02, ν_R_ = 0.03, ν_I_ = 0.2, η_LPC_ = η_FPC_ = η_OFC_ = 0.01. Time steps correspond to 10 ms approximately. Inputs are presented during 20 time steps. Model behavior (phase diagram, dynamics) is robust to reasonably large parameter variations.

In simulations we specified the task-sets, instruction cues involved in the experiment and we assigned values to the feedforward projection matrix connecting the input to reward layers in OFC and determining expected reward values, as these values were supposed to be learned during training. Unless stated, all task-sets last 200 time steps. For every simulation described below, the model performed with good accuracy, i.e. over 90% (except for simulations of episodic memory retrieval, see related section).

### Model phase diagram

Using computer simulations, we analyzed the network dynamics in different situations by varying onsets of external cues and associated expected reward values (see Fig. 2). These simulation results have been reported in detail in (Koechlin 2007). Here, we only outline the results.

**Figure 2.**
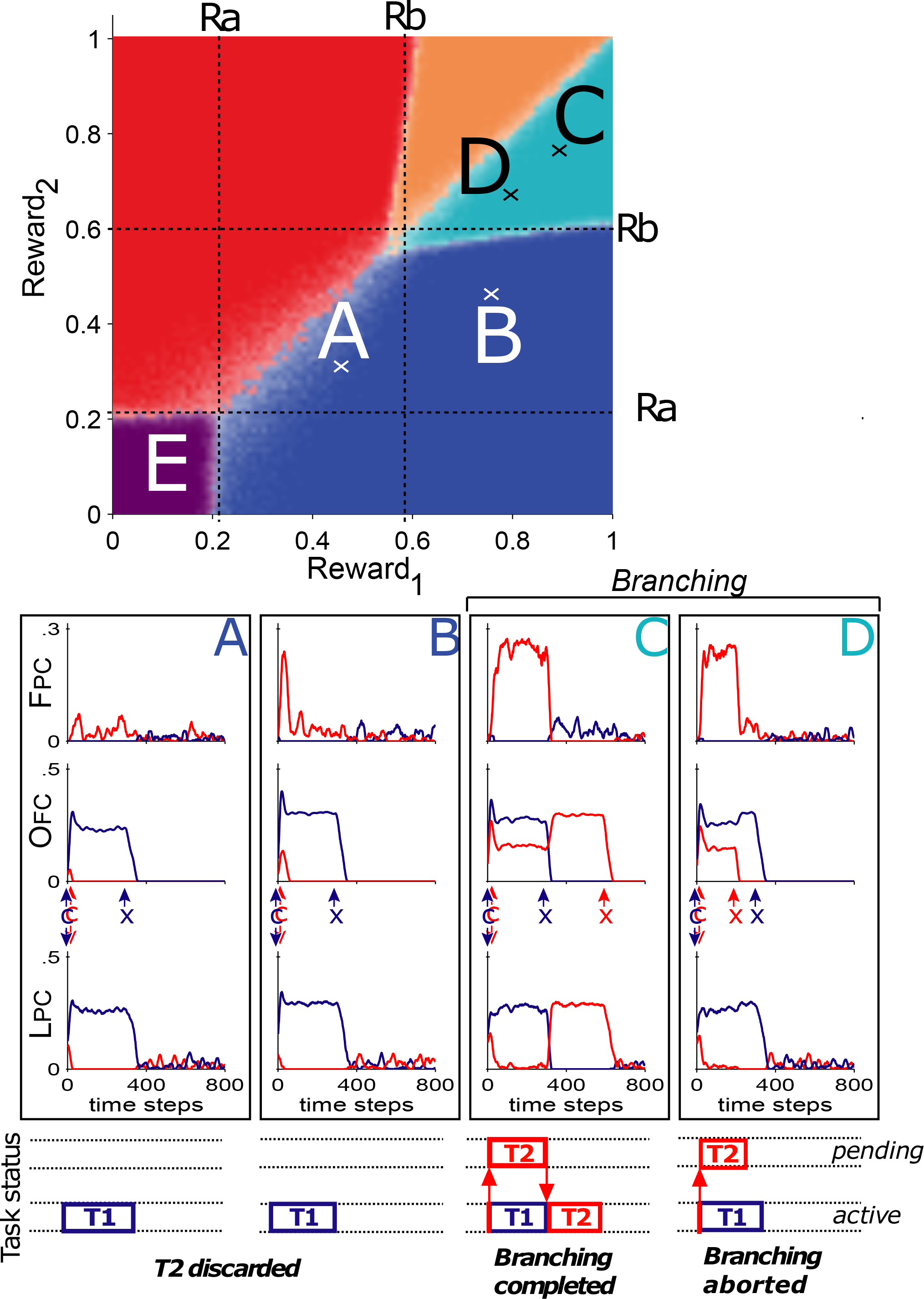
Phase diagram describing network behavior with respect to expected reward values. The phase diagram was computed from computer simulations of the network behavior in response to an external cue associated with two distinct task-sets (T1 and T2). The diagram shows five regions, delineating values of the two expected rewards (R1 and R2) where a given qualitative network behavior predominates. **A-D**: Insets provide examples of network dynamics in each region of the phase diagram. Blue and red lines represent activity of units encoding task-sets 1 and 2 respectively in LPC, OFC-reward layer-and FPC (bottom, middle and upper graphs respectively). Duration of task-set performance was set to 200 time steps; ‘x’ arrows indicate task-set completion. Purple region: discarding of both task-sets, no taskset performed. Blue region: R2 too low, only task-set 1 is executed (insets A & B); note that when R2 is close to Rb (inset B), T2 induces short phasic activations in FPC. Orange region: R1 > R2 > Rb, T1 is executed first while T2 is temporarily stacked in FPC. T2 is then retrieved following completion of T1 (inset C), unless R2 is lowered under Rb during execution of T1 (inset D). Turquoise region: R2 > R1 > Rb, T1 is executed first, then T2, branching occurs. Region boundaries are blurred as noise induced uncertainty about network outcome: paler dots around these boundaries indicate less certain behavior type. Rm is the minimum reward value for sustained activation in LPC; Rb is the minimum reward value for sustained activation in FPC, i.e. for cognitive branching to occur.

First, consider an external cue *C* cueing multiple task-sets *T1, T2, T3* …, each task-set being associated with expected reward *R1, R2, R3* …, given cue C, respectively. For clarity, we assume that *R1* > *R2* > *R3* > … The least rewarding task-sets *T3* and followings are always ignored and discarded from the system, *i.e.* the system cannot process more than the two most rewarding task-sets. Furthermore, when all expected rewards are lower than a minimal threshold *Rm*, all task-sets are ignored and the system remains at *rest.* When only the reward expected from the most rewarding task-set is larger than *Rm* (i.e. *R1 > Rm> R2)*, the most rewarding task-set *T1* is selected in LPC and guides subsequent behavior, while the others are simply ignored. When the rewards respectively related to the two most rewarding task-sets are both larger than threshold *Rm (R1 > R2 > Rm)*, the network behavior depends upon a second reward threshold *Rb*-referred to as the *branching* threshold— which is larger than threshold *Rm* (*Rb* > Rm). If the second most rewarding task-set is lower than *Rb (R2 < Rb)*, this task-set (*T2*) is ignored and only *T1* is selected in LPC for guiding subsequent behavior.

In contrast, if the second largest reward R2 is larger than *Rb*, then the associated taskset *T2* is selected and maintained in FPC in a pending state, while the most rewarding task-set *T1* is activated in LPC and guides current behavior. In other words, a branching process starts if the second most rewarding task-set is rewarding enough to be maintained in a pending state rather than discarded. This situation perpetuates until any of the two related rewards drops below the branching threshold *Rb*. If the future reward expected from pending task-set *T2* drops below *Rb*, then the pending task-set *T2* is discarded and the most rewarding task-set *T1* keeps being active in LPC and guides behavior; thus, the branching process is aborted. In contrast, if *R1* drops below *Rb*, then task-set *T1* is discarded and the pending task-set *T2* encoded in FPC automatically migrates to LPC and starts guiding subsequent behavior. The branching process is thus completed, allowing the execution of the pending task-set upon completion of the ongoing task-set.

A very similar phase diagram is observed when task-sets are successively cued (data not shown)(see Koechlin E and A Hyafil, 2007).

### Simulation results

The model was used to simulate subjects’ performance and anterior prefrontal activations in several behavioral paradigms: multitasking (Koechlin E *et al.*, 1999), cognitive integration (Christoff K *et al*, 2001), prospective memory (Burgess PW *et al.*, 2001), attentional set-shifting and episodic retrieval (Buckner RL, 2003) paradigms.

### Multitasking (Koechlin E *et al.*, 1999)

In this experimental study, stimuli were pseudorandom sequences of visually presented upper-or lower-case letters from the word ‘tablet’ (Fig. 3, bottom). In a control condition, only upper-case letters were presented and subjects had to decide whether two successively presented letters were also in immediate succession in the word ‘tablet’. In the other conditions, both lower-and upper-case letters were presented. In a delay condition, lowercases were distractors that subjects simply had to ignore while carrying on the same uppercase task as in the control condition. In a task-switching condition, subjects had to stop the current task whenever letter-case changed and to initiate a new (‘tablet’) task from the current letter (this letter being matched with ‘T’ by convention). Finally, in a branching condition, subjects had to respond to uppercases as in the delay condition and to lowercases as in the task-switching condition, so that during sequences of lowercase letters, subjects had to temporarily maintain the uppercase task in a pending state until completion of the lowercase task. Additional distinctive instruction cues were presented every 10 letters indicating subjects to switch between conditions. Experimental results revealed that, compared to the control condition, the frontopolar cortex was activated in the branching condition (only residual activity was also found in the task-switching condition). The activation in lateral prefrontal cortex was equivalent in the delay and task-switching conditions, and it was greater in the branching condition.

**Figure 3.**
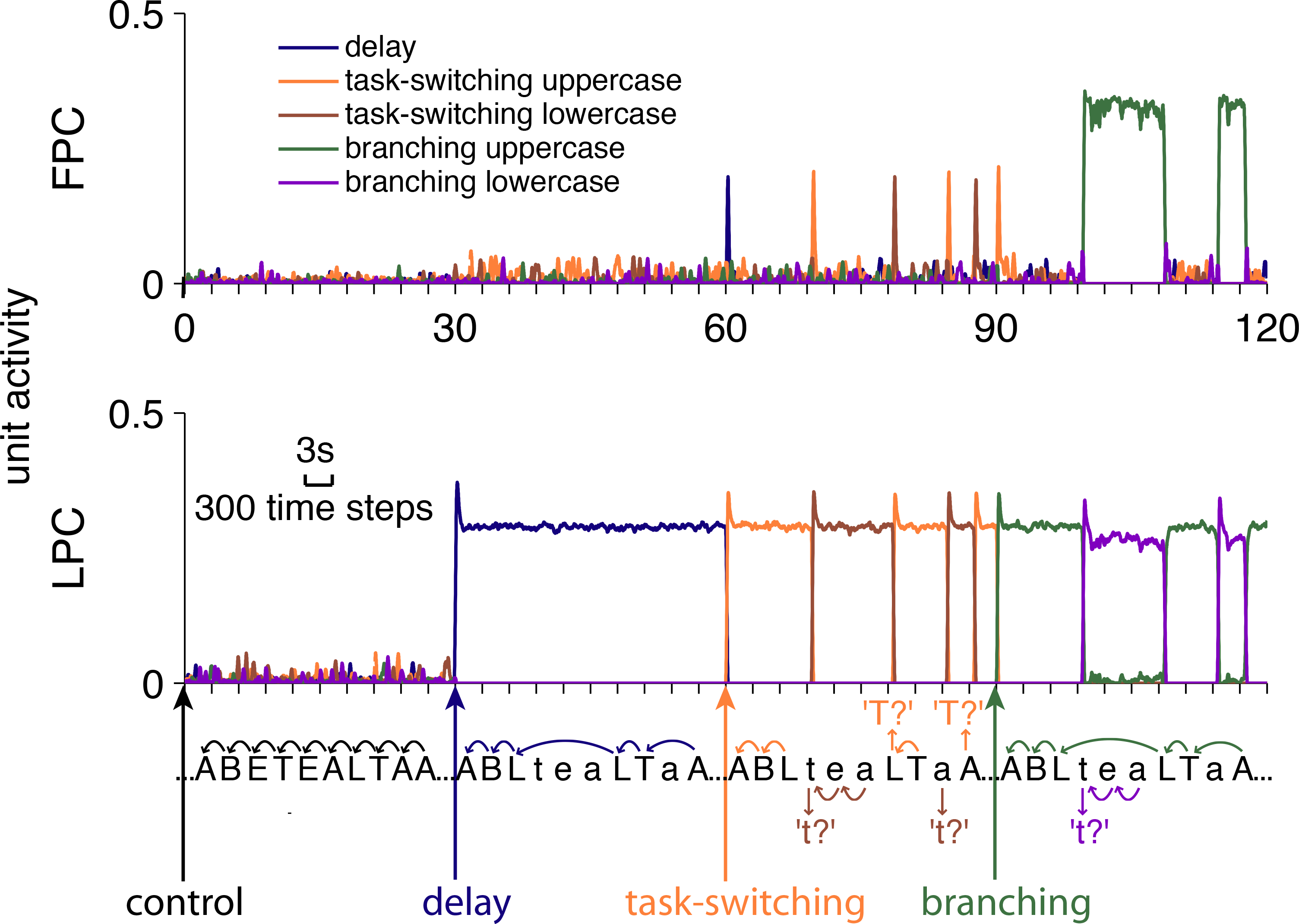
Simulation of the original ‘cognitive branching’ experiment. Graphs represent superimposed dynamics of FPC and LPC units encoding five distinct task-sets (blue, delay; orange, task-switching uppercase; brown, task-switching lowercase; green, branching uppercase; purple, branching lowercase). Letters were presented every 300 time steps (« 3 seconds) as shown by the letter series displayed at the bottom. Color arrows indicated occurrences of instruction cues (every 10 trials) related to the control, delay, dual-task, and branching conditions. Note that sustained FPC activations occurred only in the branching condition, whereas residual phasic FPC activity occurred in the task-switching condition. Rewards associated to each task-sets were set at 0.8. LPC and OFC inputs included letter cases and instruction cues.

This behavioral paradigm included *a minimum* of 5 task-sets in order to enforce behavioral rules and prevent cross-talks across conditions. One task-set specifically corresponded to the behavioral rules in the delay condition (to match two successive uppercase letters only). In contrast, task-switching and branching conditions required two independent task-sets each, one for uppercase letter decisions, the other for lowercase letter decisions. Finally the control condition did not need to be encoded as a distinct task-set, since correct performance in this condition may use behavioral rules associated with *any* other condition.

We modeled this behavioral protocol assuming that delay, task-switching, and branching instruction cues were connected to LPC units coding for the delay, uppercase task-switching and uppercase branching task-sets respectively, and to OFC units coding for associated expected rewards: expected rewards associated with each task-set was set to a fixed value in response to the corresponding instruction cues and vanished in response to other instruction cues. Additionally, LPC and OFC units coding for the lowercase task-switching and branching task-sets received excitatory inputs when lowercase letters appeared. In the task-switching condition, additionally, expected rewards associated with the task-switching uppercase task-set vanished when lowercase letters appears. Finally, because no prefrontal control was assumed to subserve cognitive processes in the control condition, no LPC units were associated with the control condition.

We simulated this model in the same way as the experimental protocol was administrated to subjects in Koechlin et al. (1999). Instruction cues related to each condition as well as lowercase and uppercase inputs were presented as successive external signals to LPC and OFC. Model activations are shown in Fig. 3.

By construction, the network in the control condition remained at rest and exhibited only background activity (this condition served as baseline only). Following delay and task-switching instruction cues, LPC units coding for associated task-sets activated. Further, in the task-switching condition, LPC activity switched back and forth between LPC units coding for uppercase and lowercase task-switching task-sets, along with changes in letter cases. In both conditions, FPC units exhibited background noise activity only, except in the task-switching condition where short phasic FPC responses were observed in task-switching trials. Following branching instruction cues, uppercase branching LPC unit activated for as long as uppercases were presented. When a lower-case letter appeared, however, LPC switched to encoding the branching lowercase task-set. Concomitantly, FPC units coding for the uppercase branching task-set activated in a sustained fashion. Then, when uppercase letters were presented again, the uppercase branching LPC unit reactivated through top-down inputs from FPC active units, while FPC activity vanished. These results grossly replicated the experimental findings reported in Koechlin et al. (1999): first, compared to the control condition, LPC activity increased in the delay condition and even more in the task-switching and branching conditions - that is additional neuronal computations occurred in LPC whenever lettercase changed (LPC was thus slightly more active in the task-switching condition in our simulations than in the original experiment). Second and most notably, FPC activity was reliable in the branching condition only, whereas only residual transient activations were observed in the task-switching condition.

### Cognitive integration

Cognitive integration requires subjects to combine the outcome of multiple tasks to produce a correct response, a cognitive process that has been associated with activations in the frontopolar cortex. For instance, in a classic integration experiment (Christoff K *et al.*, 2001), subjects performed an adapted version of the Raven Progressive Matrix tasks consisting of 3×3 matrices of visual items with the bottom right item missing (Fig. 4A). Subjects had to infer the missing item from relationships between visual features of the presented items (e.g. color, shape, or position). For example, in figure 4, the missing item of the bottom matrix is a green octagon. In each trial, the number of relationships between composing items varied. In 0-relational matrix trials, all displayed items were identical. In 1-relational trials, all items were composed of the same visual features but one that consistently varied across either matrix rows or columns. In 2-relational trials, two features consistently varied, one across rows and another across columns: only such multi-relational trials required cognitive integration. Experimental results revealed that lateral prefrontal activations increased together with subject’s response times, independently of relational complexity, whereas frontopolar activations were observed in 2-relational trials only, regardless of response times (also see (Kroger JK et al., 2002) for similar results obtained with greater number of relationships).

**Figure 4.**
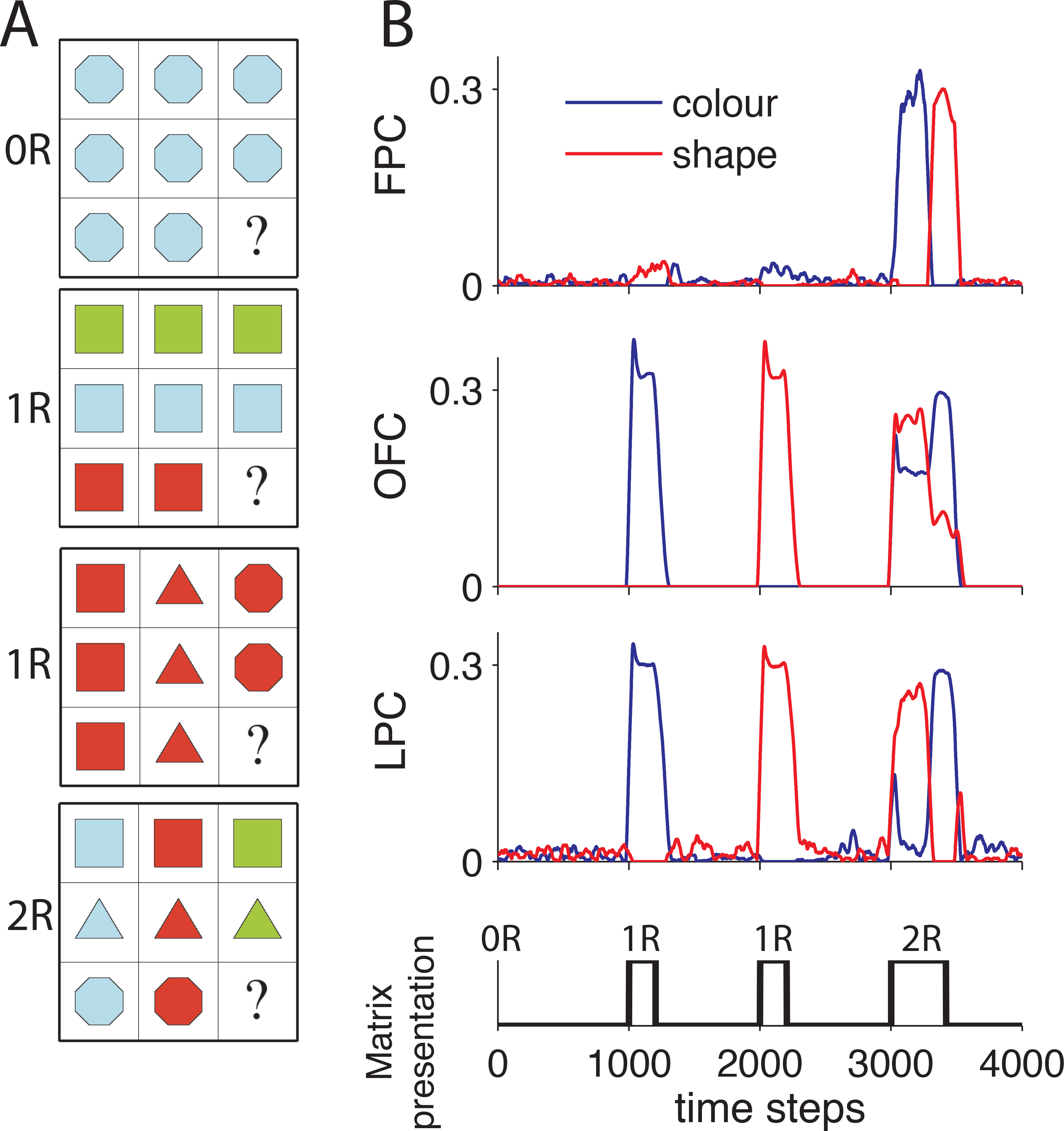
Simulations of a ‘cognitive integration’ paradigm. A. Typical examples of matrix stimuli used in the experiment. Relational complexity varies from 0-relational matrices (0R, all cells identical) to 1-relational (1R varying either only in colors, or only in shapes) and 2-relational (2R varying in both dimensions). Stimulus onsets were interspaced by 1000 time steps (10s), and stimuli offset once the network performed a response. **B**: Graphs show LPC, OFC and FPC unit activity coding for the ‘color’ and the ‘shape’ task-sets (blue and red lines respectively).

To model these tasks, we assumed that orienting attention to a specific visual feature and computing the appropriate feature for the missing item corresponded to a specific taskset. We further hypothesized that upstream perceptual areas filtered out trivial features and selected those varying across matrix items. These varying features were simply assumed to activate LPC and OFC units coding for the corresponding task-sets and related reward values. Expected reward values associated with each task-set decreased once the corresponding taskset was completed (i.e. the corresponding visual feature was computed). In addition, expected reward values related to all task-sets decreased with increased numbers of key features identified for the missing item. Finally, integration of distinct, individual features were simply assumed to result from retrieval of previously executed task-sets, by incrementally building a robust, single representation of the missing item in a perceptual space.

In 0-relational trials, LPC and FPC exhibited background noise activity since no external cue occurred (see Fig. 4B). In 1-relational trials, activity was observed only for LPC and OFC units coding for the task-set computing the non-trivial feature. This activity lasted until this feature was computed and associated expected rewards vanished. In 2-relational trials, we found that LPC units coding for the two task-sets were alternatively activated, while FPC units coding for these same task-sets activated in opposite fashion and deactivated when all expected rewards vanished. Thus, during multi-relational trials, the FPC retains the previously computed feature during computation of the others, which subsequently enables their integration into a final unified representation. The simulations replicated experimental findings (Christoff K *et al.*, 2001): first, duration of LPC activations increased from 0-to 1-and 2-relational tasks, reflecting increased amount of time required for successful inference of all features; second, FPC activated in two relational trials only, enabling the maintenance of a neuronal representation ofto recollect previously computed features while computing remaining features.

According to our simulation, the frontopolar cortex is specifically engaged during 2-relational or higher integration tasks in order to maintain and recollect previously inferred features during computation of others. Thus, contrary to the integration hypothesis, the frontopolar cortex here is not directly involved in integrating computed features per se. Integration is rather viewed as a perceptual process that gradually builds mental visual objects from individual features in posterior brain areas. The model suggests that the frontopolar cortex contributes to the reactivation and reinstatement of previously built mental objects in order to integrate new visual features. Such an account could arguably be extended to other integration paradigms like analogical reasoning tasks (Bunge SA *et al.*, 2004; Dobbins IG and S Han, 2005), which require to maintain previously computed inferences or relationships while performing a second inference task.

### Prospective memory

Using Positron Emission Tomography, (Burgess et al. 2001) addressed the issue whether FPC activations reflect either the maintenance or the execution of delayed intentions (i.e. pending tasks). Subjects first performed a baseline task on visually presented stimuli (e.g. to press a key in the direction of the larger of two displayed numbers, see Fig. 5, bottom). Then, with no prior training, subjects were instructed to proceed on this task except on certain target stimuli matching a given criteria (e.g. when both numbers were even) for which subjects had to produce an alternative response. In an *execution* condition, target stimuli appeared in 20% of trials, whereas in an *expectation* condition, there were actually no target stimuli. Experimental results revealed similar frontopolar activations in both conditions compared to baseline, and an unexpected reduction of lateral prefrontal activations in the execution compared to the expectation condition. The authors concluded that the frontopolar cortex contribution to prospective memory consists in maintaining delayed tasks in memory rather than retrieving them for execution.

**Figure 5.**
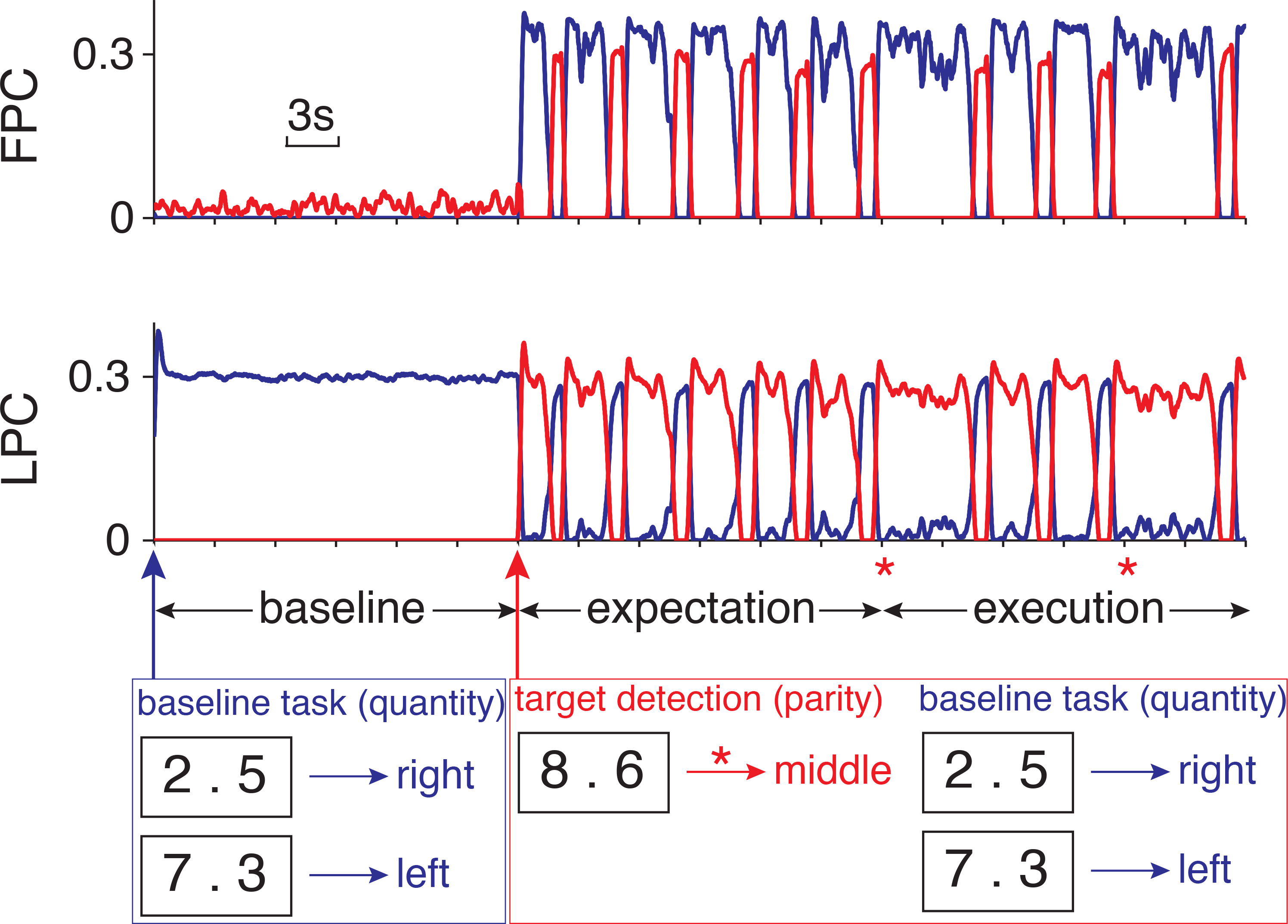
Simulation of ‘prospective memory’. We simulated only one of the four experiments described in the original study; analogies between the procedures naturally extend the validity of our results to the other experiments. Graphs show activity in LPC and FPC units corresponding to the baseline and target detection task-sets (blue and red lines respectively). Ticks on the horizontal axis represent onset of stimuli (pair of numbers) that occur every 300 time steps (i.e. 3 seconds). As in the experimental protocol, the baseline condition is presented first, corresponding to the performance of the baseline task-set alone (quantity task, blue lines). Six trials later, an external cue appears (red arrow), corresponding to the instruction to perform the additional target detection task. Target trials (red asterisks) occur only in the execution condition. Insets provide examples of stimuli with associated response (left, middle or right button press) in each condition.

Formally, this protocol required subjects to first determine whether presented stimuli were targets, then to either produce the alternative response or perform the baseline task. In principle, these two tasks - target detection (with associated response production) and baseline task - could be represented as a single task-set. However, this seems unlikely since subjects were not trained on performing these two tasks together. We thus modeled these two tasks as independent task-sets encoded in distinct LPC units. The reward scheme associated with these two task-sets was set as in the experimental protocol (Burgess PW *et al.*, 2001) and in accordance with temporal-difference models of reward evaluation. Subjects were financially rewarded only for target stimuli, when they detected them and provided the associated response. The amount of money they received depended however on their global performance on non-target trials from the baseline task. Accordingly, in our simulations, the expected reward value was higher for the target detection than baseline task-set before every trial (1 vs. 0. 65). Moreover, reward value for the baseline task remained constant over trials, while reward value for the target detection task temporarily decreased below the baseline reward level (down to 0.4) whenever a non-target stimulus was detected (since no money reward was expected in these trials). After non-target trials, the expected reward for target task-set increased back to its original value, because target-bound rewards could possibly be obtained in subsequent trials. Finally, LPC and OFC units coding for the baseline task-set received inputs signaling the onset of baseline condition, whereas units coding for the target task-set were cued by the instruction indicating how to respond to target stimuli.

Our simulations revealed that LPC encoded the baseline task-set during the baseline condition, while only background activity was observed in FPC (Fig. 5). Following instructions signaling target stimuli, LPC started to code the target detection task-set, and the baseline task-set was transferred to FPC. Then, whenever the ongoing target detection task-set detected non-target stimuli, task-sets encoded in LPC and FPC temporarily swapped as a result of transient changes in relative expected reward values: LPC started to code the baseline task-set, enabling baseline task execution, while the target detection task-set was transferred to FPC. Task-sets encoded in FPC and LPC swapped back when expected reward values reset to their original values at trial completion. Importantly, no swap occurred during target trials.

These results are in good agreement with the original PET findings (Burgess PW *et al.*, 2001) indicating that polar and lateral prefrontal regions were activated in both execution and expectation conditions compared to baseline, with maximal contrast in the frontopolar cortex. Consistently, in the model LPC exhibited sustained activity in all conditions and additional transient activations during expectation and execution conditions. In contrast, FPC exhibited both sustained and phasic activations in expectation and execution conditions only. Moreover, because phasic activations resulting from task-set swapping between LPC and FPC occurred only in non-target trials, which were more frequent in the expectation condition, the model accounts for previous findings showing greater prefrontal activations in the expectation than the execution condition (Burgess PW *et al.*, 2001).

More generally, the model accounts for polar and lateral prefrontal activations in all prospective memory paradigms (Burgess PW et al., 2003; den Ouden HE et al., 2005; Okuda J et al., 1998). According to the model, the frontopolar cortex activates in such paradigms because subjects have to manage two unrelated task-sets, i.e. the baseline (ongoing) task and the target detection or prospective memory task, one being provisionally maintained in FPC while the other is being executed. Moreover the model indicates that the frontopolar cortex starts exhibiting sustained activations when subjects receive instructions to perform an alternative response on target trials. Consistently, a previous event-related potential study of prospective memory revealed slow-wave frontopolar activations onsetting at instruction presentation in trials where targets were subsequently successfully detected (West R and K Ross-Munroe, 2002).

### Attentional set-shifting

Pollmann et al. (Pollmann et al. 2001) have reported challenging results related to the role of the frontopolar cortex, revealing its involvement in simple visual attention tasks. In their experiments, subjects viewed grids of horizontally moving green items and were instructed to press a button whenever one item (referred to as “target”) deviated in one perceptual dimension, that is whenever they saw a red square or an obliquely moving square (see Fig. 6A). The task was administered in two separate blocked conditions. In a *pop-out* condition, no additional distractors were presented, so that targets popped out from the visual display. In a *distractor* condition, little red and blue squares and little green square moving obliquely were additionally presented. Accordingly, targets were defined as the conjunction of either size & color (large & red) or size & motion (large and oblique) attributes. In both conditions, reaction times on *change* trials (when the target differed from that in the immediately preceding trial) were longer than on *repeat* trials (when the same target was repeated twice) (Fig. 6B). Such a switch cost was larger in the distractor than pop-out condition. The switch cost, however, was not observed in an additional control condition where the two possible targets varied within the same visual dimension (e.g. red squares vs. blue squares) (Pollmann S, 2001). Moreover, Lepsien et al. (Lepsien et al. 2002) found similar phasic frontopolar activations in both repeat and change trials in the distractor condition (Fig. 6D, right), which is consistent with the involvement of the frontopolar cortex in relational integration (see above). Surprisingly however, in the pop-out condition, phasic frontopolar activations were also observed on change trials (Fig. 6D, left), although this condition presumably involved automatic visual processes only. Consistently, (Pollmann et al. 2007) showed that patients with frontopolar lesions had considerable deficits in performing the distractor condition. In the pop-out condition, by contrast, frontopolar patients made very few errors, but were disproportionably slower than healthy subjects on change trials (Fig. 6C). According to the authors, these findings challenge classical theories of frontopolar function (see introduction), especially since the frontopolar cortex was activated in pop-out visual processes. Here, we addressed this issue by simulating the task using our model.

**Figure 6.**
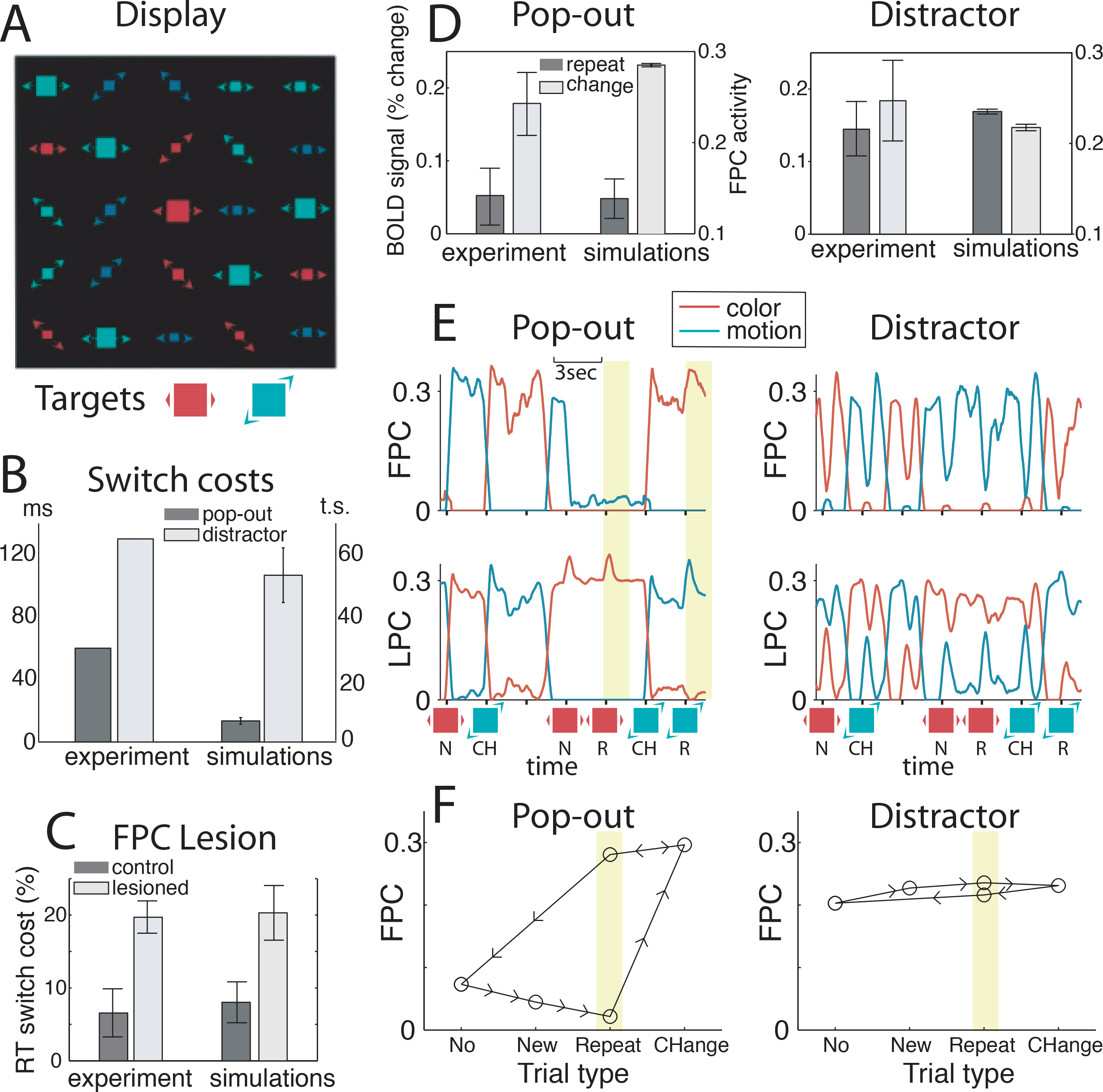
Simulation of attentional shifting. A: Illustration of a display containing a color red target in the distractor condition. Targets, present on 60% of trials, could either be a large red square moving horizontally or a large green square moving sideways. Adapted from (Weidner, 2002). **B:** Reaction times switch costs (i.e. the difference between RTs on change trials and RTs on repeat trials) in both conditions (dark bar: pop out; light bar: distractor), in the original experiment (Weidner R *et al.*, 2002) and in model simulations (right). All behavioral and activation measures reported here were obtained by averaging the results from 20 simulations of 20 trials. **C:** Effect of lesion of the FPC module (right), compared with original data on patients with lesion of lateral frontopolar cortex (left, from (Pollmann S *et a.*, 2007). Bars show relative RT switch costs (expressed as the percentage slowing of change trials compared to repeat trials), both for controls (dark bars: healthy subjects in the experiment, and full model in the simulations) and lesioned (light bars). Error bars represent 95% confidence interval for original fMRI data, and standard error across blocks for simulated data. **D:** Comparison of FPC mean activity in model simulations with the activity of the lateral frontopolar cortex in the original fMRI experiment (Weidner R *et al.*, 2002), separately for repeat trials (dark bars) and change trials (light bars), in the pop-out (left panel) and distractor (right panel) conditions. **E:** LPC and FPC unit activity in the pop-out condition (left panel) and distractor condition (right panel). Red and green lines represent activity of units related to the color and motion task-set, respectively. Ticks mark the onset of every trial; small figures denote the presence of a color or motion target for the ongoing trial. N stand for new target trials, i.e. target trials following a no-target trial; R stand for repeat trials, CH for change trials. **F:** Schematic hysteresis cycles showing how FPC activations on repeat trials in the pop-out condition depended on the recent history of preceding trials. Circles represent mean activations on a corresponding trial type, and arrows show possible ‘trajectories’ for sequence of trials. Note that such an hysteresis cycle was in the pop-out condition (left panel), but not in the distractor condition (right panel).

We simulated this attentional task using two distinct task-sets associated with the two possible targets: a first task-set was assumed to detect red (large) squares, while the second detected obliquely moving (large) squares. The key assumption was that external inputs to LPC triggering each task-set depended upon visual displays, so that input intensity associated with each task-set varied as the amount of perceptual evidence for red color (first task-set) or oblique motion (second task-set). Moreover, we assumed that the rewarding value associated with a task-set in the OFC increased in presence of perceptual evidence related to this taskset. Conversely, the rewarding value decreased only when the task-set was selected in LPC for detecting the associated target but the target was actually absent.

Simulation results showed that in the pop-out condition, task-set selection in the LPC was driven by perceptual bottom-up inputs in every target trial, so that LPC units selected the task-set associated with the target, thereby reflecting automaticity of pop-out processing (see Fig. 6E, left). As expected, this bottom-up selection process was slightly longer on change than repeat trials, so that the model exhibited small but reliable reaction time costs on change compared to repeat trials (Fig. 6B). On non target trials, LPC units simply kept on encoding the previously selected task-set. FPC units behaved in a more complex way, exhibiting hysteresis cycles (Fig. 6F, left): in repeat trials subsequent to non-target trials, no task-sets were encoded in FPC, whereas in repeat trials subsequent to change trials, the FPC encoded the non-target task-set (Fig. 6E, left). In change trials, by contrast, FPC encoded the no-target task-sets irrespective of previous trials. Such a behavior occurred because, in non-target trials, rewarding values associated with both task-sets decreased given that both task-sets were successively active in the LPC (branching occurred) but none of the targets were detected. Consequently, on the next trial, the non-target task-set remained discarded from both the LPC and FPC due to its weak rewarding value (no perceptual evidence for it), while the LPC selected the target task-set, driven by both bottom-up perceptual inputs and an increased rewarding value. The situation perpetuated until the next change trial. Then, the rewarding values of both task-sets became large again, so that target and non-target task-sets were encoded in the LPC and FPC, respectively (Fig. 6E). The FPC was then continuously activated until another non-target trial occurred. Thus, in agreement with empirical results, frontopolar activations in the pop-out condition were larger in average on change than repeat trials (Lepsien et al. 2002) (see Fig. 6D).

In the distractor condition, perceptual evidence for each task-set was large and comparable in every trial because of the numerous distractors. For the same reason, on every trial both task-sets were *a priori* associated with large rewarding values. As a result, task-set selection in LPC was not driven by bottom-up inputs. Instead, on every trial LPC kept on encoding the task-set selected in the previous trial, while the FPC encoded the other one (Fig. 6E & 6F, right). When the previously selected task-set appeared unsuccessful and its associated rewarding values decreased (for no target trials and change trials), the LPC switched to the other task-set thanks to top-down interactions from the FPC. Thus, in this condition, task-switching in the LPC was endogenously guided by changes in success/reward expectations rather than by bottom-up perceptual inputs. In accordance with empirical data (Lepsien J and S Pollmann, 2002), the model therefore exhibited a much larger switch cost in the distractor than pop-out condition (Fig. 6B). Furthermore, the model showed comparable frontopolar activations on repeat and change trials in the distractor condition (Fig. 6D, right).

Finally, we simulated a lesion of the FPC by clamping all FPC unit activity to zero, leaving the rest of the network intact. The lesioned model was unable to accurately perform the distractor condition (achieving chance level on target trials), because in the intact model task-set selection was endogenously guided by LPC-FPC interactions. In the pop-out condition, the lesioned model accurately performed the task since task-set selection was simply driven by perceptual bottom-up inputs. However, in agreement with the behavioral performance of frontopolar patients (Pollmann S *et al.*, 2007), reaction times in the lesioned model was much larger on change than repeat trials (Fig. 6C). In the intact model, such switch costs were reduced because the alternative task-set was encoded in the FPC, on change trials as well as on repeat trials subsequent to change trials. Consequently, task-set switching in the intact model was facilitated because it was driven by both perceptual bottom-up inputs and additional top-down FPC interactions in a large proportion of trials.

### Episodic memory retrieval

Episodic memory retrieval tasks, which require subjects to make judgments about past events, primarily engage the medial temporal lobes, including the hippocampus that store associations between stimuli (represented in the temporal lobe) and the context in which they occurred (spatial/temporal context etc. see review in (Frankland PW and B Bontempi, 2005; Miyashita Y, 2004)). Episodic retrieval tasks additionally recruit prefrontal regions (Buckner RL, 2003; Buckner RL and ME Wheeler, 2001; Miyashita Y, 2004). First, ventrolateral prefrontal regions are involved in effortful intentional retrieval, regardless of the success of the retrieval attempt, presumably providing *pre-retrieval* top-down signals to the hippocampus for enhancing the recollection of appropriate contextual associations (referred to as *memory cueing*)(Dobbins IG *et al.*, 2002; Fletcher PC et al., 1998; Simons JS and HJ Spiers, 2003). Second, dorsolateral prefrontal regions are engaged on successful retrieval trials, regardless of retrieval effort, seemingly reflecting *post-retrieval* processing like the selection of appropriate responses (Henson RN et al., 1999). Furthermore, frontopolar activations have been repeatedly found in episodic retrieval tasks (Fletcher PC and RN Henson, 2001; Velanova K et al., 2003). In contrast to hippocampal and lateral prefrontal regions, however, frontopolar activations in episodic memory tasks remain highly variable and difficult to interpret beyond the general agreement that it should reflect some “high-level control operations” (Simons JS and HJ Spiers, 2003).

Here, we show that the branching model accounts for the variable involvement of frontopolar regions in episodic memory retrieval tasks. For that purpose, we simply assumed that LPC units representing lateral prefrontal processing reciprocally interact with an additional processing module representing the hippocampal contribution to episodic memory (Fig. 7A). In agreement with empirical data, we assumed that distinct LPC units represent task-sets associated with pre-retrieval *(memory cueing)* and *post-retrieval* processing (response selection). Briefly, the general idea is that FPC units actively maintain representations of pre-retrieval task-sets during execution of post-retrieval task-sets, i.e. the frontopolar cortex maintains appropriate retrieval cues during processing of retrieved memories.

**Figure 7.**
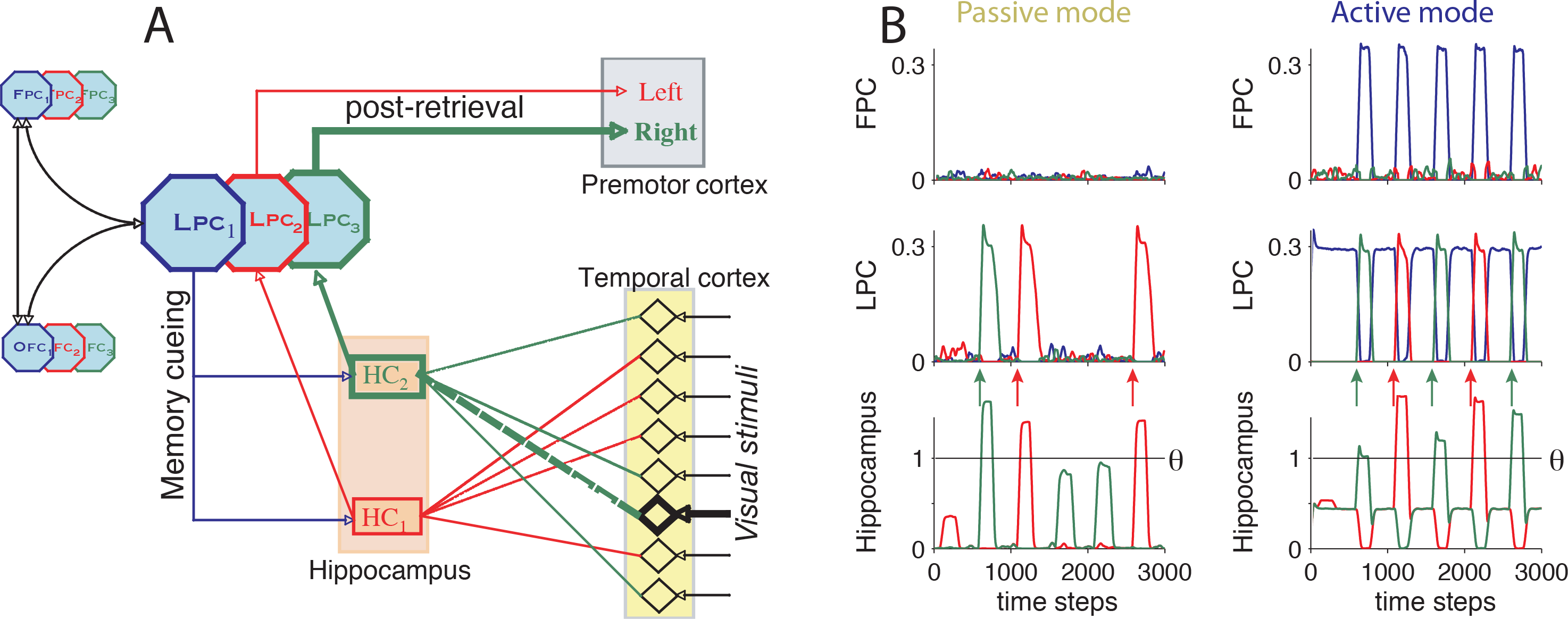
Simulation of ‘episodic retrieval’. A: The extended model including two additional modules representing neuron assemblies in the hippocampus (HC) and temporal cortex (TC). HC_1_ and HC_2_ represent the two distinct contexts in which stimuli may appear. Connection weights between the two modules vary as the depth of encoding of stimuli and associated contexts. Otherwise the model is identical to the one shown in figure 1. T1 represents the memory-cueing task-set, whereas T2 and T3 represent post-retrieval task-sets associated with distinct HC units. **B:** HC, LPC and FPC unit activity during episodic retrieval in the passive and active modes. Arrows indicate trials with successful recollection, i.e. where the HC unit activity reaches the recollection threshold θ.

We simulated a prototypical episodic memory task requiring subjects to recall which of two possible contexts (e.g. words presented in distinct lists or at different spatial positions) stimuli had been previously presented. The additional module HC representing the hippocampus coded for the contexts in which stimuli appeared (Fig. 7A). HC units received excitatory projections from additional input units (TC units) corresponding to neurons in the temporal cortex coding for stimuli (e.g. words). Connections from TC to HC units simply represented associations between stimuli and the context in which they appeared (Fig. 7A). Connection weights were set during a first encoding phase, so that each occurrence of stimulus s within context *c* strengthened the connection weight ρ_sc_ between unit TC_s_ and unit HC_c_. Thus, connection weights depended upon the number of stimulus presentations and varied as the *depth of encoding.*

During the retrieval phase, a random stimulus from the study list was presented (i.e. its corresponding TC unit was activated) every 500 time-steps. Recollection of the associated context occurred when the corresponding HC unit activated above a given “recollection” threshold θ ((Fortin NJ et al., 2004; Yonelinas AP et al., 1996)). In that case, the HC unit provided inputs to LPC and OFC units related to post-retrieval task-sets (with associated rewards worth 1). In particular, the associated LPC unit represented the *post-retrieval* task-set that selects the appropriate response given the recollected context (e.g. ‘left’ if the word was presented in list 1, ‘right’ otherwise). If no HC units reached “recollection” threshold θ following stimulus presentation, then recollection failed and a random response was generated (Yonelinas AP *et al.*, 1996).

First, we separately considered two distinct modes of retrieval, namely the *active* and *passive* mode reflecting two opposite extremes of retrieval effort (Buckner RL, W Koutstaal, DL Schacter, AD Wagner et al., 1998). In the active mode, the *memory-cueing* task-set was associated with a large rewarding value (R = 0.8) through OFC units, so that LPC units coding for the *memory-cueing* task-set were activated at the beginning of the test phase. HC units were then activated through both top-down excitation from these LPC units and bottom-up inputs from TC units, so that HC units often activated above recollection threshold θ. In the *passive* mode, by contrast, the rewarding value associated with memory cueing was weak (R = 0) so that there was no top-down excitation from LPC to HC units, and recollection relied only upon bottom-up inputs from TC to HC units. HC units had a linear response to their input:

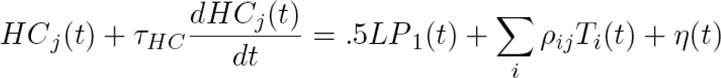

with τ_Hp_=10, LP_1_ being activity of the memory cueing task-set.

Simulation results showed that in both modes, model performance improved with increased depth of encoding (Fig. 8A). Furthermore, after shallow encoding the model performed much better in the active than passive retrieval mode (78% vs. 52% correct responses for stimuli presented only once during the encoding phase), whereas after deep encoding, both modes yielded similarly high accuracy (99 vs. 97% for stimuli presented four times). In the passive mode, as expected, only LPC units coding for the post-retrieval task-sets were activated, whenever the associated HC units activity reached recollection threshold (Fig. 7B, left). No activity above background level was observed in FPC. In the active mode (Fig. 7B, right), memory-cueing LPC units were tonically activated during the retrieval phase, except after successful recollection, which triggered activations of post-retrieval LPC units. On such occasions, FPC temporarily encoded the memory-cueing task-set until completion of the post-retrieval task. The branching mechanism then enabled automatic reactivation of memory-cueing LPC units for subsequent trials. Thus, FPC was only recruited in successful recollection trials during the active mode. This result is consistent with event-related imaging studies showing frontopolar activations in successful compared to unsuccessful recollection trials (Henson RN *et al.*, 1999; Ranganath C et al., 2000; Rugg MD et al., 2003; Simons JS et al., 2005; Wheeler ME and RL Buckner, 2003). Moreover, in our simulation, FPC activated only during post-retrieval processing, which is consistent with empirical data showing long-latency frontopolar activations (Buckner RL, W Koutstaal, DL Schacter, AM Dale et al., 1998; Reynolds JR et al., 2005).

**Figure 8.**
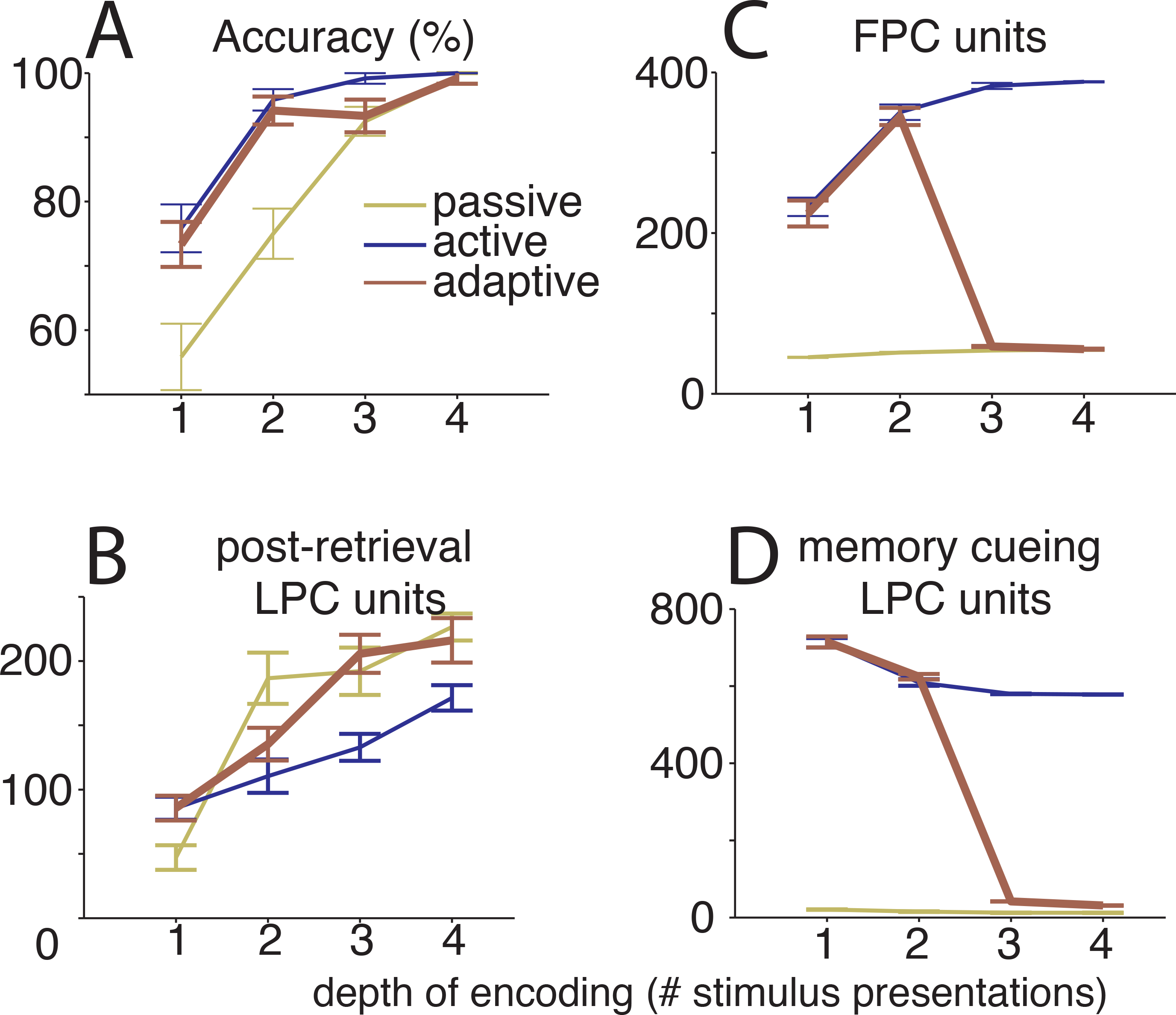
Behavioral and activation measures from model simulations of an episodic retrieval experiment, in passive (green lines), active (blue lines) or adaptive mode (brown lines), as a function of depth of encoding (measured by the number of stimulus presentation during the encoding phase). **A.** Model accuracy. **B.** Activations of the LPC units related to the postretrieval task-sets (T2 & T3). **C.** Mean activations of the FPC units. **D.** Activation of the LPC unit related to the memory cueing task-set (T1). All activation data points represent total activity in the unit population averaged over trials. Errors bars are standard error across 20 blocks of 6 trials.

We then studied transitions between the active and passive modes in relation to depths of encoding. Presumably, the active mode is enhanced, provided that it significantly improves recollection performance compared to the passive mode, which only occurred after shallow encoding (Buckner RL, W Koutstaal, DL Schacter, AD Wagner *et al.*, 1998). Accordingly, we assumed that the rewarding value of memory-cueing was proportional to (i.e. 3 times) the accuracy gain in the active compared to the passive mode, which was computed for various depths of encoding from the simulations described above. Simulations were then run in this more realistic *adaptive* mode.

As expected, we observed that in the adaptive mode too, model performance improved with increased depths of encoding (Fig. 8A). For low encoding depths, model accuracy corresponded to that in the active mode, whereas for large encoding depths, model accuracy was similar to that in the passive model. In parallel, activations of post-retrieval LPC units also increased with depth of encoding (Fig. 8B). By contrast, activations of memory-cueing LPC units showed the opposite pattern and gradually decreased with increased depth of encoding (Fig. 8D), reflecting a gradual transition from the active to the passive mode. Finally, FPC activity varied as a bell-shaped function of depth of encoding (Fig. 8C): FPC activity remained weak when encoding was either very shallow or very deep, but strongly increased at intermediate encoding depths. This bell-shaped curve was observed because FPC activations jointly depended on two factors that varied in opposite ways with depth of encoding. On one hand, FPC activated only on successful recollection trials whose frequency increased with depth of encoding. On the other hand, FPC activity was contingent upon activations of memory-cueing LPC units, which decreased with increased depth of encoding. Thus, at shallow encoding, mean FPC activity was weak because successful recollection was rare, even if the model performed in the active mode, whereas at deep encoding, FPC was poorly activated because the model performed in the passive mode. These bell-shaped variations of FPC activity may explain apparent discrepancies in frontopolar activations observed across episodic retrieval experiments: (Buckner et al. 1998) reported increased frontopolar activations during deep compared to shallow encoding, whereas the reverse result was observed in a more recent study (Velanova K *et al.*, 2003). In agreement with the present model, however, maximal frontopolar activations in both studies were found in conditions corresponding to an intermediate depth of encoding (two word presentations in the encoding phase, retrieval accuracy: ˜85%), where the model precisely exhibited maximal FPC activity (Fig. 8C).

### Discussion

In this study, we examined the hypothesis that cognitive branching, i.e. the process underlying our ability to postpone the execution of a primary task until completion of another, is the core function of the frontopolar cortex that explains its engagement in a variety of cognitive paradigms. For that purpose, we used a neurocomputational model of cognitive branching for simulating the engagement of anterior prefrontal regions in such cognitive paradigms (Koechlin E and A Hyafil, 2007). The model shows how branching processes may dynamically emerge from functional interactions between three processing modules representing polar, lateral and medial/orbital prefrontal regions. It suggests that the frontopolar cortex works as a stack by maintaining a representation of a (highly rewarding) pending task, while lateral prefrontal regions are encoding the ongoing task associated with a larger rewarding value. This situation perpetuates until ongoing task becomes less valuable than the pending task, in which case the pending task migrates to lateral prefrontal regions and starts being active. More generally, this model exhibits several behaviors depending upon rewarding values of both tasks (see Koechlin E and A Hyafil, 2007 for a detailed discussion). While with the parameters used here only one task-set at a time could be maintained in a pending state, this pending capacity could easily be amplified simply by lowering the level of inhibition in frontopolar cortex. The model could thus how two task-sets can simultaneously be encoded in frontopolar cortex (Donoso M *et al.*, 2014). Moreover, simulation results show that the model accounts for frontopolar activations observed in multitasking paradigms that explicitly involve branching processes. In agreement with empirical data (Braver TS and SR Bongiolatti, 2002; Koechlin E *et al.*, 1999; Sakai K and RE Passingham, 2003), the model exhibited sustained frontopolar activity when a task was postponed until completion of another, whereas the frontopolar cortex was not recruited when a task was simply delayed for future execution. Moreover, the model exhibited residual phasic frontopolar activity during simple task-switching, which accounts for incidental empirical findings of frontopolar activations in task-switching paradigms (e.g. Dreher, 2003). The model also correctly predicts that the frontopolar cortex activates during exploratory behaviour, allowing to maintain counterfactual choices and constantly update their reward value when new evidence is provided (Daw N *et al.*, 2006; Boorman E *et al.*, 2009, 2011).

Previous brain imaging studies revealed that the frontopolar cortex is also involved in relational reasoning, when subjects have to combine the outcomes of multiple tasks to produce appropriate responses (Christoff K *et al.*, 2001; Kroger JK *et al.*, 2002). Accordingly, cognitive integration, i.e. the integration of outcomes from multiple cognitive tasks, has been proposed as a basic frontopolar function (Christoff K *et al*, 2001; Ramnani N and AM Owen, 2004). Our simulations however show that, rather than the final integration of outcomes *per se*, frontopolar activations in such paradigms may alternatively support the maintenance of outcomes from previously executed tasks during performance of subsequent tasks. This branching account for frontopolar activations in integration paradigms assumes that single cognitive tasks are performed successively before their integration. This is consistent with previous behavioral results showing that multiple tasks can only be processed serially through central/prefrontal executive function (Dux PE et al., 2006; Pashler H, 2000). Frontopolar activations were also reported in more complex reasoning and problem-solving tasks, such as the Tower of London task (Baker SC et al., 1996; Christoff K and JD Gabrieli, 2000). Cognitive branching *a fortiori* accounts for these results, given that such complex tasks require subjects to devise and evaluate several sub-tasks while keeping in mind superordinate plans. Consistently, no frontopolar activations were found in the Tower of London task when subjects directly performed a sequence of moves, without any reasoning on alternative plans (Owen AM et al., 1996). Frontopolar activations were also reported in much simpler tasks which clearly involved cognitive branching such as a counting task requiring subjects to count the number of successively presented stimuli matching a given criteria (MacLeod AK et al., 1998), i.e. to continuously switch between the main counting task and the stimulus categorization task (in the absence of any external switch cue).

Frontopolar activations were also observed in prospective memory paradigms (subjects, while performing a given baseline task on series of stimuli, were additionally instructed to perform a different task in response to specific target stimuli), even when no target stimuli appeared (Burgess PW *et al.*, 2001). Based on this result, the frontopolar cortex was assumed to simply implement the maintenance of delayed intentions during the performance of another task, even when the intentions were actually never executed. Our results suggest, however, that this prospective memory paradigm implicitly involves cognitive branching, because subjects had to switch without any explicit cue between two independent tasks, namely the detection of target stimuli and the baseline task. Indeed, this switching process requires subjects to maintain one task in a pending state while performing the other one. Here the two tasks could not be integrated together into a single fixed sequence of tasks, which would have spared the need for cognitive branching, since it would have required extensive training for building such sequences. With such training, in contrast, we predict that the involvement of cognitive branching and consequently frontopolar activations would decrease. Consistently, (Burgess et al. 2003) found that frontopolar activations decreased with practice in prospective memory paradigms. Thus, the model accounts for the disengagement of the frontopolar cortex in prospective memory when delayed intentions are linked to the ongoing task. In summary, the branching hypothesis states that the frontopolar cortex is involved in prospective memory, provided that delayed intentions are expected to be valuable enough and have not been linked to the ongoing task or any external cue, such as remembering to buy bread for dinner when passing by the bakery (McDaniel MA and GO Einstein, 2000).

One difficulty in interpreting neuroimaging data is to disentangle between activations associated with processes genuinely required to perform a task and those corresponding to additional processes reflecting a particular strategy used by subjects to improve performance. For example, frontopolar activations have been often, although unsystematically, reported in several task-switching and attentional set-shifting paradigms, where task selection is driven by external inputs (Braver TS et al., 2003; Pollmann S, 2001). In the model, switching between task-sets does not require the engagement of the frontopolar cortex. However, model simulations revealed that maintaining the subsequent task-set in a pending state in FPC speeds up subsequent task-switching in LPC (see attentional-set shifting section in Results), because FPC units provide additional selective inputs to LPC units. The recruitment of FPC units occurs however only when task-sets are rewarding enough. Thus, the model predicts that the frontopolar cortex activates in task-switching paradigms only when fast responses and accuracy are especially emphasized. Consistently, patients with lateral frontopolar lesions were found to accurately perform a pop-out attentional-set shifting task but with much greater switch costs than normal subjects (Pollmann S *et al.*, 2007). By contrast, despite nearly normal performance on classical intelligence tests, frontopolar patients are dramatically impaired in multitasking paradigms requiring to manage multiple concurrent tasks and to switch in and out multiple rewarding tasks in absence of any external cues (Burgess PW, 2000). In agreement, the model necessitates an intact FPC module to switch in and out multiple tasks without external cues. Thus, the model accounts for both neuroimaging and neuropsychological data and explains why the frontopolar cortex may be strategically engaged in simple attention and task switching protocols, while being mandatory in multitasking behaviors. In particular, the model predicts that frontopolar activations obey hysteresis cycles in attentional set-shifting paradigms developed by (Pollmann et al. 2001; 2007), so that the frontopolar cortex is “strategically” activated according to the recent history of task trials rather than to specific task requirements in each trial.

Frontopolar activations have been reported in a number of episodic memory retrieval paradigms and especially in source-memory protocols requiring subjects to retrieve the context in which stimuli appeared (Ranganath C *et al.*, 2000; Rugg MD *et al*, 2003; Suzuki M et al., 2002). However, no robust predictors of frontopolar activations in episodic retrieval tasks have been established so far. The present model suggests that FPC is engaged in such episodic retrieval paradigms in order to improve recollection performance by efficiently combining two distinct task-sets: cueing memories of contexts in which stimuli were presented and processing products of retrieval attempt, which are respectively implemented in ventrolateral and dorsolateral prefrontal regions (Buckner RL and ME Wheeler, 2001; Dobbins IG *et al.*, 2002; Fletcher PC *et al.*, 1998). Thus, retrieval accuracy is improved by the involvement of FPC in maintaining the memory-cueing process in a pending state during the execution of post-retrieval subtasks, which enables the resumption of memory cueing processes on subsequent trials despite the absence of external cues. As shown in Results, the model accounts for the frontopolar activations observed in previous episodic memory retrieval studies. In particular, our model predicts a bell-shape relationship between the depth of stimulus encoding and frontopolar activity, which reconciles apparently inconsistent empirical findings (Buckner et al. 1998; Velanova K *et al.*, 2003).

Cognitive branching enables to retrieve and perform a postponed task, even in absence of any cues that can trigger its execution. Conversely, we predict that cognitive branching will not occur when such reminder cues are expected, thus sparing the cognitive cost associated with the engagement of frontopolar regions. For instance, when two tasks are repeatedly performed in succession, sequential associations between the two tasks may gradually develop, providing internal associative inputs triggering the execution of the second task after completion of the first one, without engagement of cognitive branching. Indeed, frontopolar deactivations have been found in various experiments during the course of learning (Burgess PW *et al.*, 2003), when such sequential associations could develop. In several studies with explicit learning of motor and task sequences (Jenkins IH et al., 1994; Koechlin E et al., 2002; Muller RA et al., 2002), the frontopolar cortex has also been reported to activate in the early learning phase, then to gradually disengage as learning progressed. We did not simulate these learning paradigms, because such simulations require elaborating additional models of taskset execution and adaptive learning, which are out of the scope of the present study. Nevertheless, the branching hypothesis accounts for the early engagement of frontopolar regions in learning, when subjects critically switch back and forth between alternative behavioral options/task-sets (exploration) drawing upon feedback to progressively eliminate irrelevant choices in search of optimal behavior. Then, when contingency or uncertainty over potential task-sets decreases, e.g. when serial associative links develop among several task-sets because of training or simply when the number of possible task-sets shrinks, the need for branching processes decreases. Consistently, previous studies showed gradual frontopolar disengagements as contingency and uncertainty over multiple task-sets decrease during learning (Yoshida W and S Ishii, 2006), exploration (Daw ND et al., 2006) or multitasking (Koechlin E *et al.*, 2000).

In summary, the present model provides theoretical evidence that multitasking, relational reasoning, prospective memory and learning explicitly or implicitly rely on cognitive branching between multiple mental tasks. Furthermore, the model suggests that the frontopolar cortex is not required but may be strategically involved in attentional set-shifting and episodic memory retrieval tasks because cognitive branching may improve performance especially in series of trials challenging fast responses and high accuracy. These results support the hypothesis that cognitive branching is the core function of the frontopolar cortex, which explains its involvement in such a variety of mental tasks. The hypothesis is further supported by recent studies showing substantial overlaps between frontopolar activations observed in these diverse cognitive paradigms (Dobbins IG and S Han, 2005; Gilbert SJ et al., 2006).

As described above, the model makes several key predictions regarding the pattern of frontopolar activations that should be observed in all these mental tasks. The model makes additional predictions regarding functional interactions between lateral, orbital/medial and polar prefrontal regions as well as between reward processing and frontopolar function. First of all, the model predicts that neuronal projections between lateral, orbital/medial and polar prefrontal regions are all excitatory, except the projection from lateral prefrontal to frontopolar regions. This latter projection from lateral prefrontal regions is predicted to exert an inhibitory influence on neuronal processing in the frontopolar cortex, i.e. to mainly target inhibitory interneurons in frontopolar regions (Barbas H et al., 2002; Barbas H et al., 2005). Nevertheless, despite this inhibitory projection, frontopolar activations only occur with concomitant lateral prefrontal activations thanks to the indirect excitatory pathways through orbital/medial prefrontal regions. Second, cognitive branching would occur between two task-sets, and consequently frontopolar activations should be observed provided that expected rewards associated with *both* task-sets are larger than a given threshold (Rb), which is greater than the reward value threshold (Rm) triggering the execution of a task-set alone. In that case, the least rewarding task-set would be postponed and encoded as a pending task-set in the frontopolar cortex, while the most rewarding task-set would be encoded in lateral prefrontal regions for governing ongoing behavior. Conversely, if one expected reward becomes lower than the branching threshold Rb, the associated task-set would be abandoned and discarded, cognitive branching would be aborted and the frontopolar cortex would disengage (in particular, the pending task-set reward may drop below threshold Rb due to occurrences of external feedbacks/events).

In conclusion, the model only focuses on neuronal processes in the frontopolar cortex and their interactions with neighboring prefrontal regions, including the orbital/medial and lateral prefrontal regions. Consequently, the model is based on an oversimplification of neuronal processes implemented in lateral, orbital/medial prefrontal regions. The model only hypothesizes that lateral prefrontal regions select task-sets based on external signals and represent ongoing task-sets. This assumption is consistent with previous neuroimaging studies showing that selection and maintenance of task-sets that govern ongoing behavior involve lateral prefrontal regions (Braver TS *et al.*, 2003; Koechlin E *et al.*, 2003). The model, however, does not account for complex neuronal processing in more posterior frontal and parietal/temporal regions involved in task-set performance and acquisition. Similarly, the model only assumes that orbital/medial prefrontal regions compute and represent expected future rewards associated with task-sets. Again, this assumption is consistent with recent empirical findings (Matsumoto K and K Tanaka, 2004; Rolls ET, 2004), although the present model certainly offers a rather simplistic view of reward processing in these regions. Finally, the model explains the activation of the frontopolar cortex as depending only on the 'pending' status of a task-set, while it was more recently found that the frontopolar cortex also monitors the reliability of the task-sets (Donoso *et al.*, 2014). This notion could be added in a more refined implementation of the model. Despite these important limitations, we believe that the model captures several key features of frontopolar function, which do not critically depend upon such simplifications. In particular, the model makes several key basic predictions that can be empirically tested to provide new insights about neuronal mechanisms underlying higher cognition.

